# EFFECT OF ORGANIC AND CHEMICAL FERTILIZERS ON THE NUTRITIONAL COMPOSITION OF *Amaranthus spinosus*

**DOI:** 10.1101/2022.06.02.494619

**Authors:** Deepa Raman, Namita Joshi, Pawan Singh Rana

## Abstract

Organic and chemical fertilizers are used in agriculture as they improve the fertility of soil and the growth of plants. So, the present investigation was designed to determine the effect of different fertilizers on the nutritional value of *Amaranthus spinosus* plant. The quantity of different fertilizers was used as 500 kg h^-1^. Some selected parameters of soil such as temperature (29.5^0^ C), moisture (3.5 %), Water holding capacity (74%), pH (8.17), organic carbon % (1.90) and organic matter % (3.28) were measured and analysed. There was no significant variation in these parameters under different fertilizers treatment. The phytochemicals of Organic fertilizers treated plant sample had higher concentration of Alkaloid (1.06), flavonoid (0.89), tannin (1.48) and saponin (1.67), ash (7.75), moisture (87.6) and protein (9.1), Fe (0.054), Zn (0.008), Cu (0.001), Na (0.0080), Ca (113) and Mg (93) when compared to the other fertilizer treatments. Inorganic fertilizer did not show any significant effect on plant growth, while the combination of organic and chemical fertilizers was found to have a significant effect. The results reveal that organic fertilizers or their combination with chemical fertilizer are better for the crops than chemical fertilizers.

## INTRODUCTION

The thorny *Amaranthus spinosus* belongs to family Amaranthaceae. It is a coarse herb with tiny green flowers (Costea *et* al, 2001) and used as a vegetable, fodder and in medicine (Ayethan *et* al, 1995). It is a good source of vitamin A, C, folic acid, thiamine, niacin and riboflavin and also contains a significant amount of minerals viz. calcium, iron, potassium, zinc, copper and manganese and the source of lysine; essential amino acid, which is obtained from Amaranthus seeds (Pee *et* al, 1995). The crop is mainly grown in India, Sri Lankaand warm temperate region of Asia from Japan to Indonesia. In India, *Amaranthus spinosus* are used to treat the bronchitis, internal bleeding, diarrhoea, excessive menstruation, blood disease, burning, leprosy and piles. It is also used as anti-inflammatory, antioxidant and hepatoprotective herbal product. (Srivastva *et* al, 2011). The grains of *Amaranthus spinosus* has high nutritious value, used in daily diet in the form of cereals, soup, cookies and snacks (Martha and Shimelis, 2012).

Organic fertilizers (cow dung manure and poultry farmyard) have a good quality of NPK which improves soil texture and yielding of crops. Organic and inorganic fertilizers are necessary for the plant growth, which provides the nutrients needed for the optimum growth of the plants. Most of the countries have been using the organic fertilizers from ancient time, whereas the use of chemical fertilizers is relatively recent due to demand of the food for rapidly growing populations at the global level. People prefer to use the chemical fertilizers as because they are easy to use and are quickly absorbed and utilized by the crops (Erisman *et*, al 2008).

The organically grown vegetables and fruits contain a good amount of minerals, vitamins and protein (Citak *et* al, 2010). Organic fertilizers also play an important role to improve the chemical and physical properties of soil. On the other side, chemical fertilizers are mainly applied for the rapid growth of plant and high yielding of a product (Stewart *et* al, 2005). But the excessive use of chemical fertilizer affects the chemical and physical properties of soil, resulted in the deficiency of nutrients from plant product (Singh *et* al, 2001). The balanced use of both types of fertilizers is essential for the growth of crop with good quality and high yields.

The main objective of this study was to understand the effect of organic fertilizers and chemical fertilizers on the nutritional composition of *Amaranthus spinosus*.

## MATERIAL AND METHODS

The present study was carried out on the cropland of village Tisang, District Muzaffarnagar, Uttar Pradesh. The cropland was cleared manually and soil parameters were recorded before applying the fertilizers. The organic fertilizer (cow dung manure: 500 kg. ha^-1^), chemical fertilizer (DAP + Potash: 500 kg. ha^-1^) and a combination of both fertilizers, organic and chemical: 500 kg. ha^-1^ i.e. T_1_, T_2_ and T_3_ respectively were applied to the soil and mixed thoroughly before sowing the seed. The unit plot size was 3×3.5 m^2^ =10.5m^2^. Seedlings were raised at a spacing of 10cm to 20cm and depth of 3cm, one seedling per hole. When plants were grown up to a height 60 cm, 20 matured plants were randomly uprooted. Moisture content, ash, protein, chlorophyll and carotenoids were estimated on green leaves of the study plant. Leaves and stem were oven dried at 74^0^ C for 24 hours and grounded using mortar-pestle.

### Physico-chemical parameters of the soil

Soil temperature was recorded by using a soil thermometer. Moisture content was determined after oven drying the fresh soil sample at 105^0^ C and expressed as a percentage of the weight of the soil sample. The Water holding capacity was determined by the method as described in Trivedy and Goel, (1998). pH was measured by using a digital pH meter. Organic carbon was determined by wet digestion method of Walkley and Black, (1934).

### Determination of Phytochemicals

Phytochemicals are naturally occurring chemical compounds in plants. Quantitative analysis of phytochemicals in the plant powder was carried out using standard methods. Alkaloids were determined by Harbone, (1973) method, flavonoid by Boham and Kocipal, (1994) method and Tannin by van-Burden and Robinsonmethod, (1981). Saponins were quantified by using the protocol described by Obadoni and Chuka, (2001).

### Determination of Proximate composition

The Proximate composition of *Amaranthus spinosus* was done in fresh green leaves. Protein was estimated by Lowry *et al*., (1951) method. Moisture content was determined by oven drying method and Ash was determined by Association of Official Analytical Chemists, (1999) method.

### Determination of minerals

Plant powder was used for the determination of Calcium and Magnesium by the titrimetric method according to Trivedy and Goel, (1998). Wet digestion sample was performed to analysis of minerals. Zinc (Zn), Copper (Cu), Iron (Fe) and Sodium (Na) were determined according to AOAC, (1999).

### Determination of Phyto-physicological attributes

Chlorophyll from the fresh plant sample was estimated by Arnon method, (1949). Different concentrations of chlorophyll were calculated by applying the formula given by Machlachan and Zalik, (1963). Carotenoid was determined using the method of Duxbury and Yentsch, (1956).

## RESULTS

The present study revealed that effect of organic fertilizer was higher on plant samples in the comparison of chemical and combination of both fertilizers.

**Figure 1:**
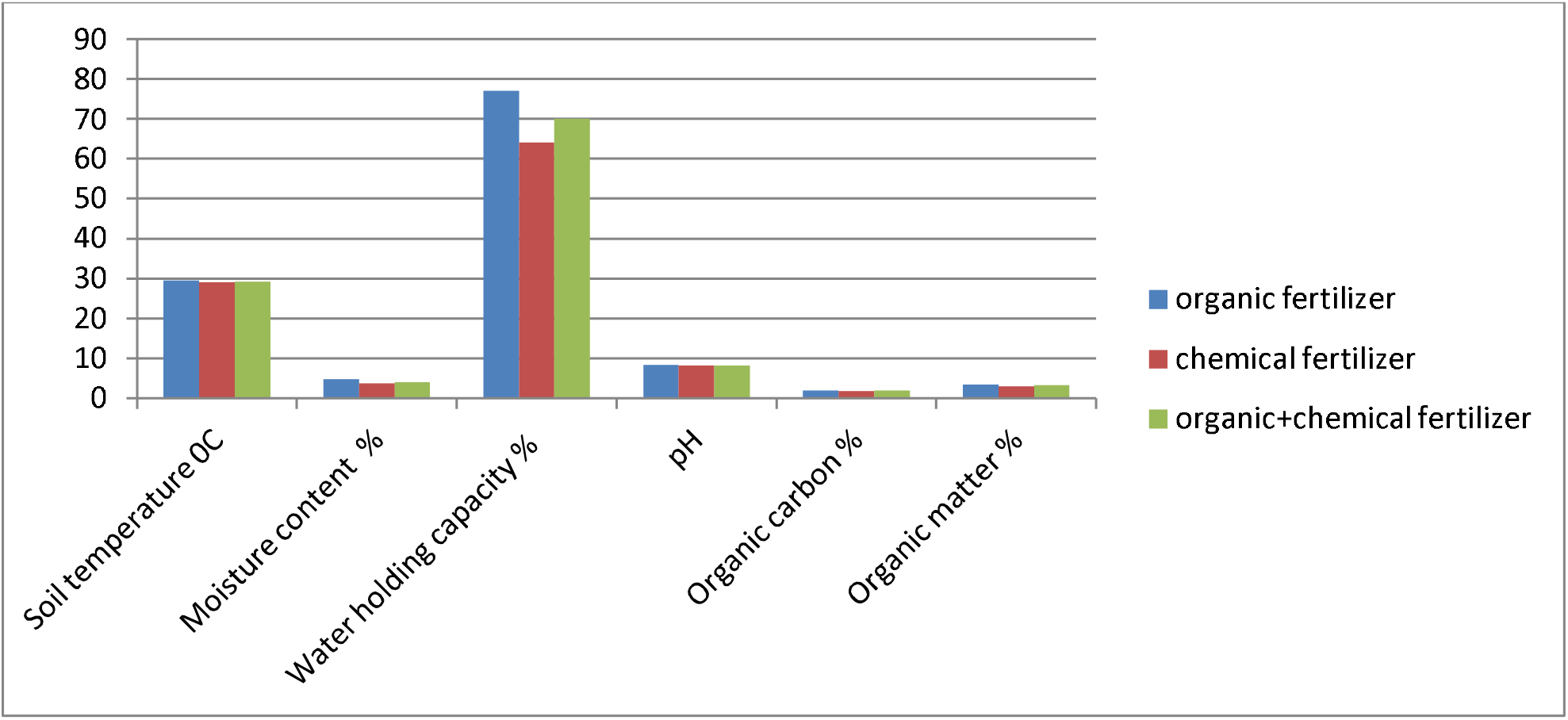
Effect of various treatment of fertilizers on soil profile.

**Figure 2:**
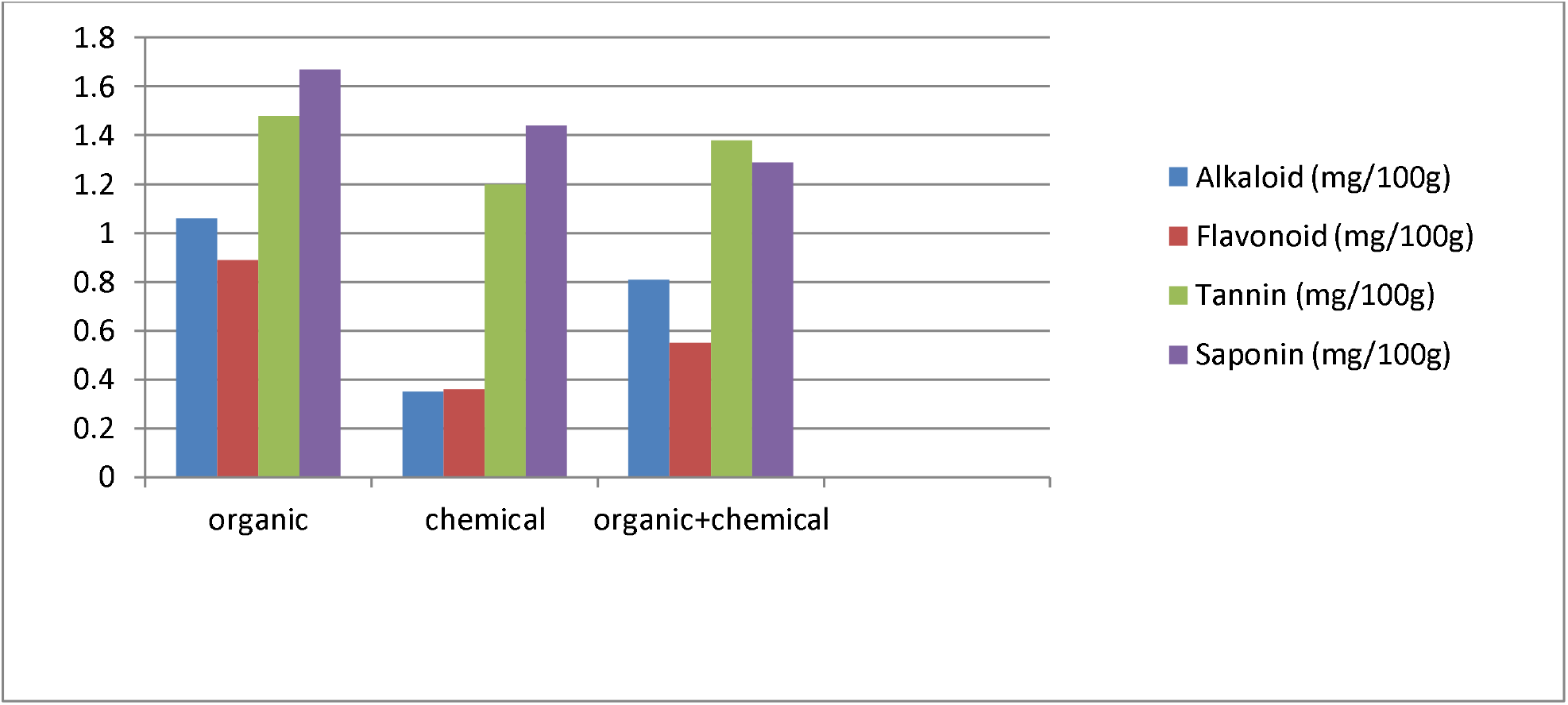
Effect of various treatments of fertilizers on phytochemicals.

Phytochemicals of *Amaranthus spinosus* was significantly (p<0.01) influenced by different treatments of fertilizers.

**Figure 3:**
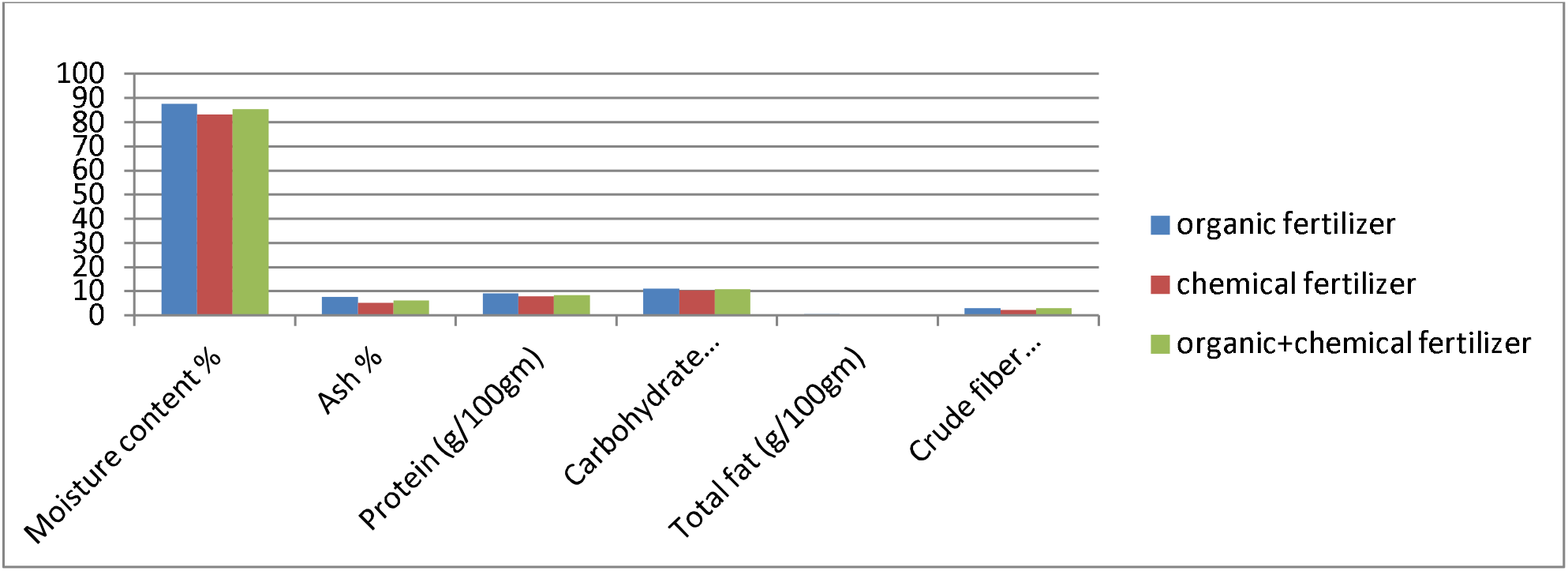
Effect of various treatments of fertilizers on the proximate composition.

Proximate composition of Amaranthus spinosus was significantly (p<0.01) influenced by treatment of organic and chemical fertilizers. The result revealed that organic fertilizer treated plant have a higher amount of proximate composition in terms of moisture, ash, protein, carbohydrate, total fat and crude fiber because organic fertilizers has a positive effect on the nutritive value of plants in comparison of chemical and combination of both fertilizers.

**Figure 4:**
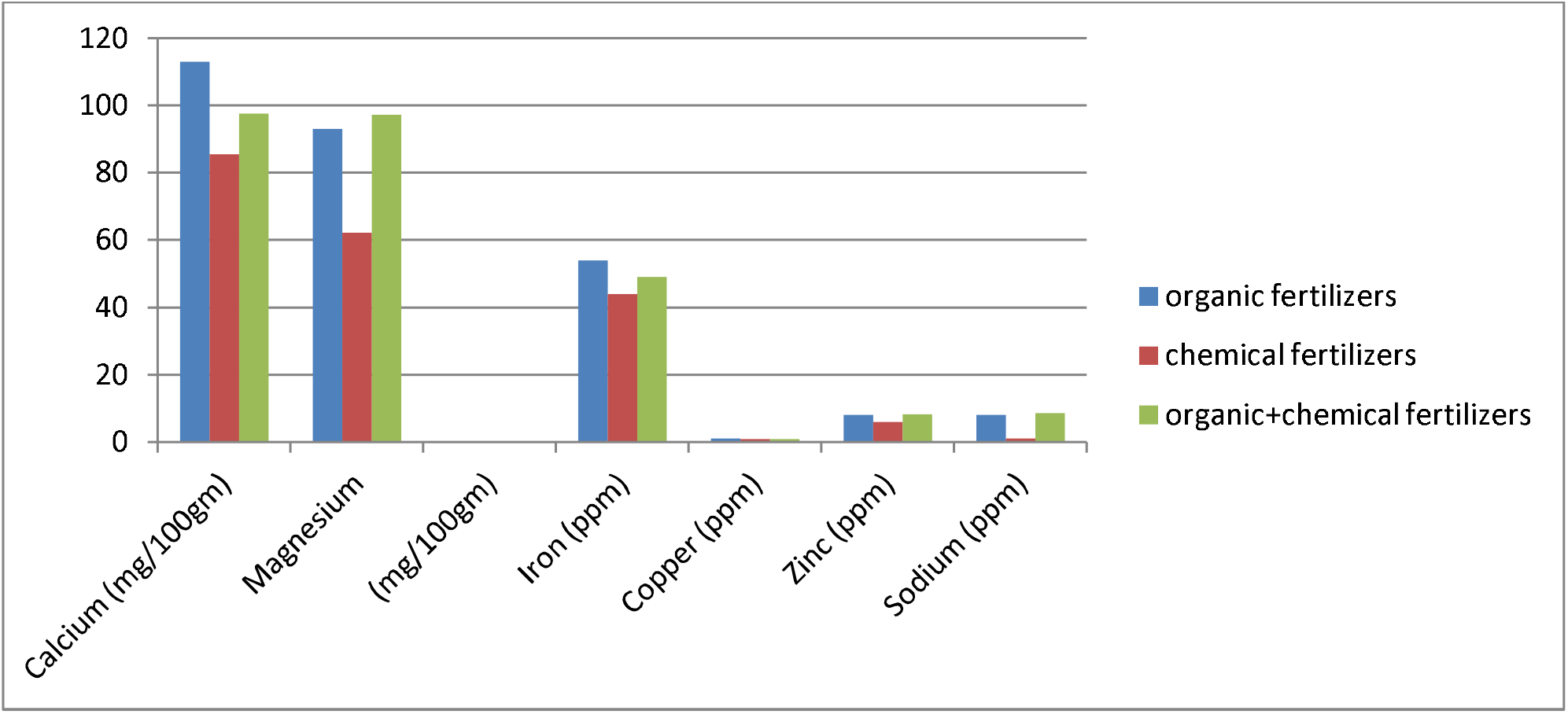
Effect of various treatments of fertilizer on minerals.

It is evident from Table 4 that Minerals in Amaranthus spinosus was significantly (p<0.04) influenced by the different treatment of fertilizers.

**Table 1:** Physico-chemical parameters of the soil under different treatments of organic and chemical fertilizers.

| S.No | Treatments<br>Parameters | T<sub>1</sub> | T<sub>2</sub> | T<sub>3</sub> |
| --- | --- | --- | --- | --- |
| 1. | Soil temperature °C | 29.5±0.09 | 29.1±0.04 | 29.2±0.04 |
| 2. | Moisture content % | 4.8±0.04 | 3.7±0.09 | 4.1±0.06 |
| 3. | Water holding capacity % | 77±0.54 | 64±0.9 | 70±0.55 |
| 4. | pH | 8.38±0.004 | 8.19±0.004 | 8.22±0.003 |
| 5. | Organic carbon % | 1.97±0.02 | 1.77±0.03 | 1.93±0.02 |
| 6. | Organic matter % | 3.40±0.05 | 3.05±0.06 | 3.32±0.05 |

(Values are mean±SE)

**Table 2:** Values of Phytochemicals of *A. spinosus* under different treatments of fertilizers.

| S.No | Treatments<br>Parameters | T<sub>1</sub> | T<sub>2</sub> | T<sub>3</sub> |
| --- | --- | --- | --- | --- |
| 1. | Alkaloid (mg/100g) | 1.06±0.03 | 0.35±0.10 | 0.81±0.12 |
| 2. | Flavonoid (mg/100g) | 0.89±0.01 | 0.36±0.02 | 0.55±0.01 |
| 3. | Tannin (mg/100g) | 1.48±0.012 | 1.20±0.023 | 1.38±0.028 |
| 4. | Saponin (mg/100g) | 1.67±0.045 | 1.44±0.062 | 1.29±0.038 |

(Values are mean±SE)

**Table 3:** Values of proximate composition of *A. spinosus* under different treatments.

| S.No | Treatments<br>Parameters | T<sub>1</sub> | T<sub>2</sub> | T<sub>3</sub> |
| --- | --- | --- | --- | --- |
| 1. | Moisture content % | 87.6 ±0.52 | 83.2±0.43 | 85.4±0.19 |
| 2. | Ash % | 7.75± 0.29 | 5.08±0.32 | 6.25±0.55 |
| 3. | Protein (g/100gm) | 9.1±0.46 | 7.9± 0.36 | 8.4±0.32 |
| 4. | Carbohydrate (g/100gm) | 11.10±0.25 | 10.41±0.28 | 10.90±0.34 |
| 5. | Total fat (g/100gm) | 0.49±0.42 | 0.27±0.22 | 0.38±0.48 |
| 6. | Crude fiber (g/100gm) | 3.06±0.37 | 2.31±0.32 | 2.89±0.28 |

Values are mean±SE

**Table 4:**
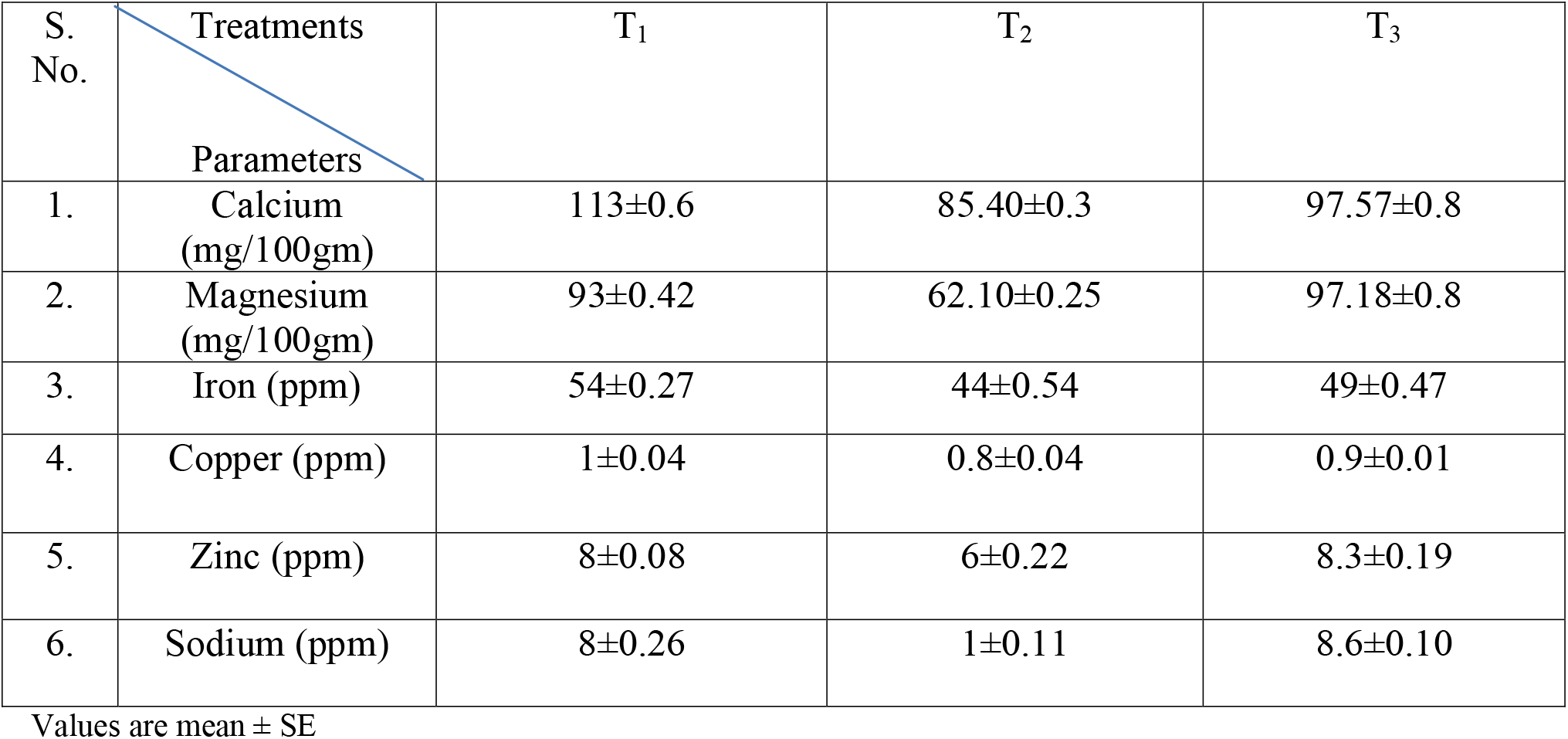
Values of selected plant mineral of *A. spinosus* under different treatments of fertilizers.

| S. No. | Treatments<br>Parameters | T<sub>1</sub> | T<sub>2</sub> | T<sub>3</sub> |
| --- | --- | --- | --- | --- |
| 1. | Calcium (mg/100gm) | 113±0.6 | 85.40±0.3 | 97.57±0.8 |
| 2. | Magnesium (mg/100gm) | 93±0.42 | 62.10±0.25 | 97.18±0.8 |
| 3. | Iron (ppm) | 54±0.27 | 44±0.54 | 49±0.47 |
| 4. | Copper (ppm) | 1±0.04 | 0.8±0.04 | 0.9±0.01 |
| 5. | Zinc (ppm) | 8±0.08 | 6±0.22 | 8.3±0.19 |
| 6. | Sodium (ppm) | 8±0.26 | 1±0.11 | 8.6±0.10 |

Values are mean ± SE

**Figure 5:**
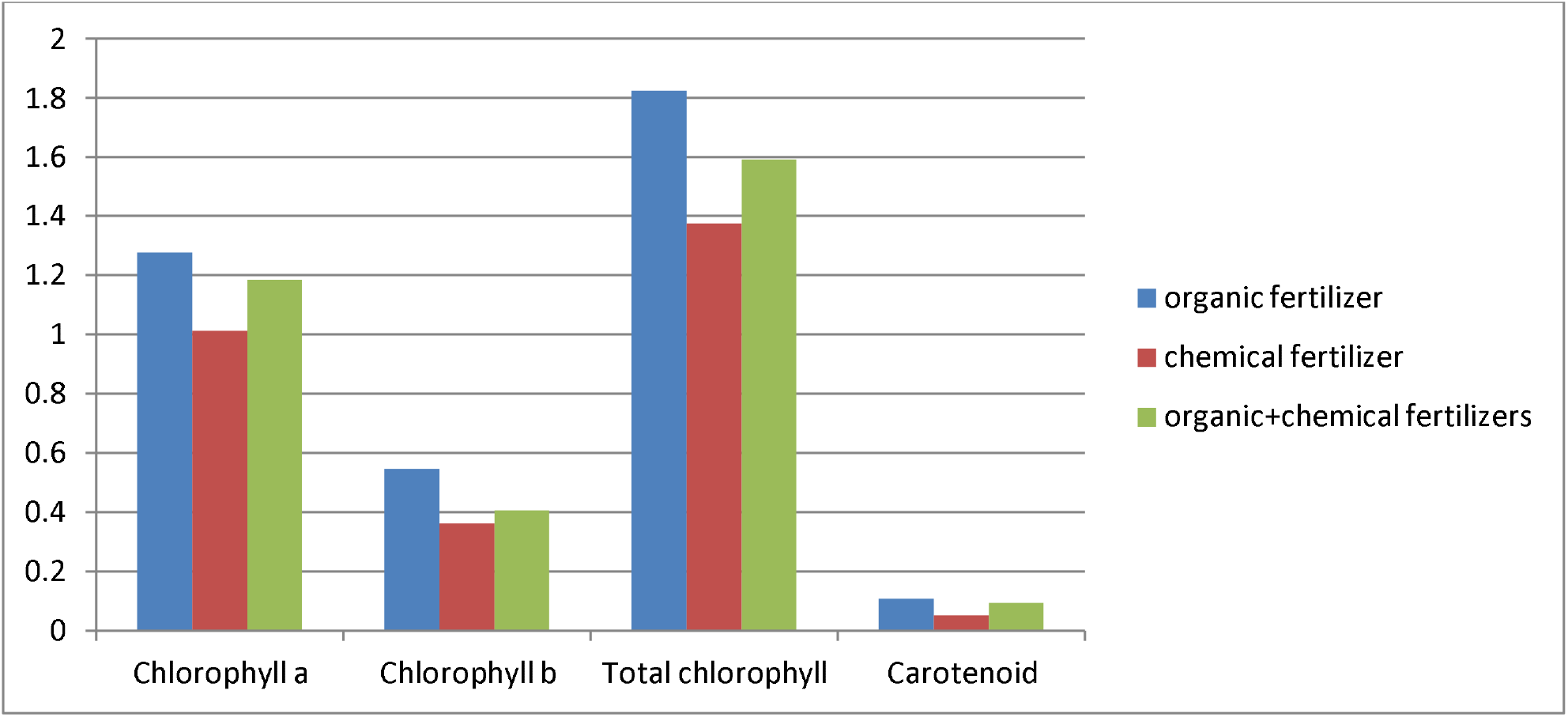
Effect of various treatments of fertilizers on phyto-physicological attributes.

In Table-5, phyto-physicological attributes of plant sample was significantly (p<0.02) influenced by the different treatments of fertilizers1.591±0.002 mg g^-1^f.w.) and carotenoid(0.094±0.002mg g^-1^f.w.).

**Table 5:**
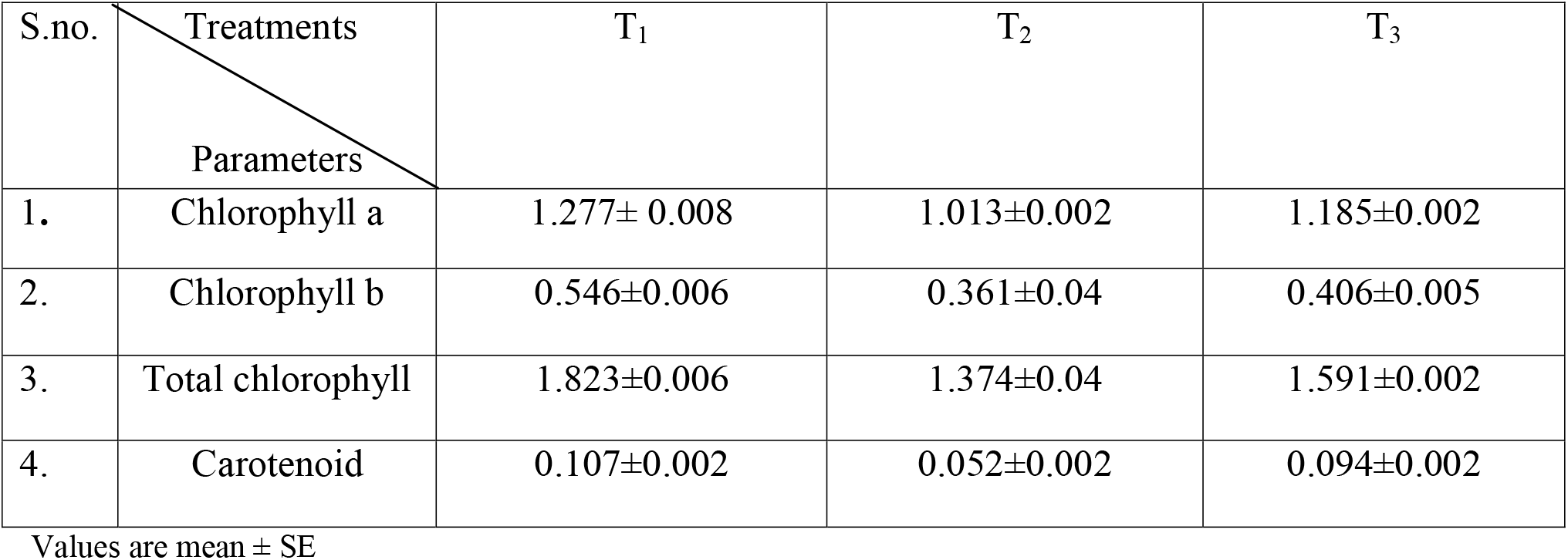
Values of few selected Phyto-physicological attributes of *A. spinosus* under different treatments (mg g^-1^f.w)

## DISCUSSION

The results revealed that organic fertilizers produced a higher amount of nutrients in comparison to the treatments of fertilizers. In the present study, highest pH was recorded in the organic fertilizer treated soil in comparison to the other treatments of fertilizers. Krishna *et* al., (2004) studied that organic fertilizers have ability to transfer the electrochemical properties of acidic soil to improve the base retention and increasing soil pH. The present study revealed that highest amount of organic matter and organic carbon was observed in the organic fertilizer treated soil in comparison to chemical and combination of both the fertilizers. Ashworth and Alloway, (2008) reported that organic matter increases with the increase of soil pH because high pH have ability to increase negative charge on the soil surface and repels negative charge in the soil solution because of this phenomena organic matter increases. The results of present study are in the agreement of Hossain and Ryu, (2017) who found that organic matter and pH of organic fertilizer treated soil was high in the comparison of chemical fertilizer treated soil sample. Organic fertilizers gradually release nutrients into soil which is responsible for the improvement of water holding capacity, pH and organic matter of soil (Suge *et al*., 2011).

In the present study, organic fertilizer treatment to the plant *Amaranthus spinosus* is responsible for the higher production of moisture content, ash, protein, carbohydrate, fat and fiber while minimum amount of mentioned parameters were observed in chemical fertilizers treatment because organic fertilizers consist of balanced nutritive elements in comparison to chemical fertilizers. Talib *et al*., (1994) suggested that organic fertilizers increase the leaves size due to level of photosynthesis increases. Amount of protein, carbohydrate and crude fiber increased due to enhancement of physiological activities. Blandino and Rezneri, (2009) suggested that nitrogen positively influence the leaf area and chlorophyll concentration which is responsible for the high yield of protein content. The present study is in the agreement of Elsheikh and Elzidany, (1996) and Makinde *et* al. (2010), who observed the highest value of proximate composition in the organic fertilizer treated plant, when compared with chemical fertilizers treated plant sample.

In the present study, organic fertilizer treated plant samples produced higher amounts of phytochemicals in comparison of chemical and combination of both fertilizers. Michael *et al*, (2010) reported that increased value of phytochemicals could depend on the availability of organic fertilizers, which can be easily decomposed and absorbed by the plants while, NPK could be lost from chemical fertilizer by leaching. The results are in accordance with the study of Coolong *et al*, (2004), Adekayode and Ogunkoya, (2011) who reported that organic fertilizers produced high concentration of phytochemicals and nutritional composition in the comparison of chemical fertilizers. The results are in accordance with the study of Mofunanya *et al*, (2014) who reported the high amount of proximate composition and phytochemicals in *Amaranthus spinosus* were in organic fertilizer treatment in comparison to chemical fertilizers treatment.

The results of present investigation revealed that concentration of calcium, magnesium, Copper, zinc, iron and sodium were higher in organic fertilizer treated plant sample in comparison to chemical fertilizer. Khadra *et al*. (2000) suggested that the availability of N, P and K was higher in the compost treatment which might be attributed for the fastest rate of decomposition and mineralization other than inorganic fertilizers. Organic fertilizers consist of balance quantity of the micronutrients which are absorbed by the plant and increase the level of nutrients. Results of the present study are in agreement with the study done by Katherine, (2007) who observed high amount of flavonoids, iron and zinc in organically synthesized food other than non-organically synthesized food. The results of the present study are in accordance of Chavan *et al*, (2015) who observed the organic fertilizers are responsible for the higher production of minerals in comparison to chemical fertilizers. Arisha *et al*. (2003) reported that organic fertilizers may stimulate the plant growth and plant nutrients than the chemical fertilizers. In the comparative study, results revealed that the high amount of Chlorophyll content and carotenoid observed in the organic fertilizers treatment while in the chemical fertilizer treated plant sample least amount of chlorophyll and carotenoid was observed. Siavoshi and Laware, (2013) reported that organic fertilizers provide macro and micro elements to the plant, which utilized for the metabolic activity to synthesize chlorophyll content. Suntoro, (2002) reported that magnesium is responsible for the chlorophyll synthesis whereas iron is responsible for the maintenance and synthesis of chlorophyll. Calcium and copper are responsible for proper functioning of the iron.

## CONCLUSION

This study dealt with the use of organic, chemical and combination of both fertilizers applied at the rate of 500 kg. ha^-1^. Organic fertilizers provide good nourishment to the plant in comparison to other fertilizers. In some cases where chemical fertilizers become a necessity to complete the demand of high yield, the farmers can opt for a suitable combination of organic and chemical fertilizers instead of chemical fertilizers. As, comparative study of different ratio of fertilizers treatment, plant showed that organic fertilizer and combination of both fertilizers treated plant sample produced a higher concentration of nutrients when compared to alone treatment of chemical fertilizers. Hence, the outcome of the study is clear in the favor of organic fertilizers and the integration of organic and chemical fertilizers to enhance the nutritive values of plant.

## Notes

### Competing Interest Statement

The authors have declared no competing interest.

